# Complete assembly of parental haplotypes with trio binning

**DOI:** 10.1101/271486

**Authors:** Sergey Koren, Arang Rhie, Brian P. Walenz, Alexander T. Dilthey, Derek M. Bickhart, Sarah B. Kingan, Stefan Hiendleder, John L. Williams, Timothy P. L. Smith, Adam M. Phillippy

**Author notes:** These authors contributed equally to this work (Sergey Koren, Arang Rhie).

## Abstract

Reference genome projects have historically selected inbred individuals to minimize heterozygosity and simplify assembly. We challenge this dogma and present a new approach designed specifically for heterozygous genomes. “Trio binning” uses short reads from two parental genomes to partition long reads from an offspring into haplotype-specific sets prior to assembly. Each haplotype is then assembled independently, resulting in a complete diploid reconstruction. On a benchmark human trio, this method achieved high accuracy and recovered complex structural variants missed by alternative approaches. To demonstrate its effectiveness on a heterozygous genome, we sequenced an F1 cross between cattle subspecies *Bos taurus taurus* and *Bos taurus indicus*, and completely assembled both parental haplotypes with NG50 haplotig sizes >20 Mbp and 99.998% accuracy, surpassing the quality of current cattle reference genomes. We propose trio binning as a new best practice for diploid genome assembly that will enable new studies of haplotype variation and inheritance.

## Introduction

Accurate reference genomes are essential for most genomic analyses, but DNA sequencing technologies cannot yet read entire chromosomes in a single pass. Instead, the genome must be reconstructed from many shorter sequences in a complex process known as assembly ^1^. Repetitive sequences longer than the sequencing read lengths prevent a complete reconstruction of chromosomes, so the assembly process typically results in a collection of contiguous sequences (contigs) that are interrupted by repeats or lack of sequencing coverage. The recent emergence of long-read sequencing technologies has dramatically improved the quality of genome assemblies by resolving many such repeats ^2^. However, even these technologies have not yet overcome the challenge of completely assembling both haplotypes of a diploid genome. Instead, most genome assembly tools simply co-assemble the haplotypes into a mosaic consensus, resulting in an assembly that does not accurately represent either original haplotype. For example, the first human shotgun assembly was built in this manner ^3^. It has since been shown that “collapsing” haplotypes into a single consensus representation introduces false variants not present in either haplotype, leading to annotation and analysis errors ^4^. Ideally, a genome should be represented as a complete set of haplotypes rather than an artificial mixture.

A common approach to the diploid assembly problem has been to skirt the issue by selecting an inbred individual to reduce the problem of haplotype variation (e.g. fly ^5^, mouse ^6^). However, this is impractical for many species and, even when possible, can result in a genome that is not representative of variation found within the natural population. An alternative approach is to use haploid, clone-based genomic libraries, as was done for the human genome project ^7^. More recently, a diploid human assembly was constructed using tiled fosmids ^8^. However, reliance on cloning is often impractical. Other attempts have been made to separate haplotypes *de novo* from whole-genome sequencing. For example, the highly polymorphic sea squirt *Ciona savignyi* was first assembled using modifications to the Arachne assembler ^9^ designed to split haplotypes based on read overlap information ^10^. However, this was an extreme case as the reference individual had an estimated heterozygosity of 4.6%. Early attempts to assemble a diploid human genome, with heterozygosity of just 0.1%, first collapsed the haplotypes into a combined assembly and then phased alleles over a short range using pairs of heterozygous variants observed on a single read or read pair ^11^. Current phasing tools operate similarly, and map sequencing reads to a reference sequence to infer blocks of variants that originate from the same haplotype ^12–14^. More sophisticated library preparations such as chromosome sorting ^15^, Strand-seq ^16^, and Hi-C ^17^ can link variants over a much longer range, delivering chromosome-scale phase blocks. However, methods that rely on reference mapping typically fail in regions of high heterozygosity and/or significant structural variation between haplotypes, yielding a limited view of genetic diversity.

A more comprehensive solution to the diploid assembly problem is to integrate haplotype separation into the assembly process itself. However, this approach is limited by the fragment length of the sequencing process, and sequencing reads alone do not always contain enough information to link variants across longer regions of homozygosity, resulting in relatively short phase blocks. As a compromise, diploid assemblers such as FALCON-Unzip ^18^ and Supernova ^19^ output “pseudo-haplotypes” that represent a single allele at each position, but do not preserve phase across long homozygous alleles or assembly gaps. In addition, these assemblers can confuse repeats with diverged alleles, leading to artifactual duplications or deletions. One potential solution is to combine long-read sequencing with additional types of information such as linked reads and/or bacterial artificial chromosomes (BACs) ^20^, Strand-seq ^21^, or Hi-C ^22^. However, no *de novo* assemblers currently integrate these data types, so this can be a manual and expensive process.

We provide a simple and cost-effective solution to the diploid assembly problem that assembles accurate, genome-scale haplotypes *de novo*. Unlike other methods that are limited to phasing individual chromosomes, we produce two complete, haploid genomes — one for each parental haplotype. Key to our method is the separation of haplotypes *prior* to assembly using a father-mother-offspring trio. Each haplotype is then assembled separately without the interference of inter-haplotype variation. Trios have long been used in genomics to infer inheritance, including for the HapMap project ^23^, the 1000 Genomes Project ^24^, and the creation of “platinum” variant catalogs ^25^. Trios were also used by *trio-sga* to simplify heterozygous diploid genome assembly ^26^, but reliance on short-read sequencing limited the haplotype-specific contigs (haplotigs) to an average size of a few kilobases. In contrast, our long-read method enables the assembly of multi-megabase haplotigs and complete parental haplotypes.

Here we introduce trio binning, and demonstrate that it reconstructs accurate and complete parental haplotypes for a wide range of zygosity and genome sizes. We first report results for benchmark datasets with both high (Arabidopsis) and low (human) levels of heterozygosity, and illustrate that prior methods do not completely recover both haplotypes of a diploid genome. We then report a novel, complete diploid assembly of an F1 hybrid between *Bos taurus taurus* and *Bos taurus indicus*, and demonstrate that the quality of each haplotype exceeds that of even the best livestock reference genomes. The closing discussion explores limitations and applications of this technique, and argues that trio binning of outbred genomes should become the new gold standard for assembling diploid reference genomes.

## Results

### Complete haplotype assembly with TrioCanu

We have implemented trio binning and haplotype assembly as the TrioCanu module of the Canu assembler ^27^. The method requires moderate coverage of short, high-quality sequencing reads (e.g. 30× Illumina) from two parental genomes to identify short, length *k* subsequences (*k*-mers) that are specific to each parent. These *k*-mers are presumed to be specific to the corresponding haplotypes of the offspring. Next, long reads are collected from an offspring of the parents to sufficiently cover both haplotypes (e.g. 80× PacBio, 40× per haplotype). Long reads from the offspring are then binned into paternal and maternal groups based on the presence of the haplotype-specific *k*-mers, and assembled separately (**Figure 1**, **Methods**).

**Figure 1.**
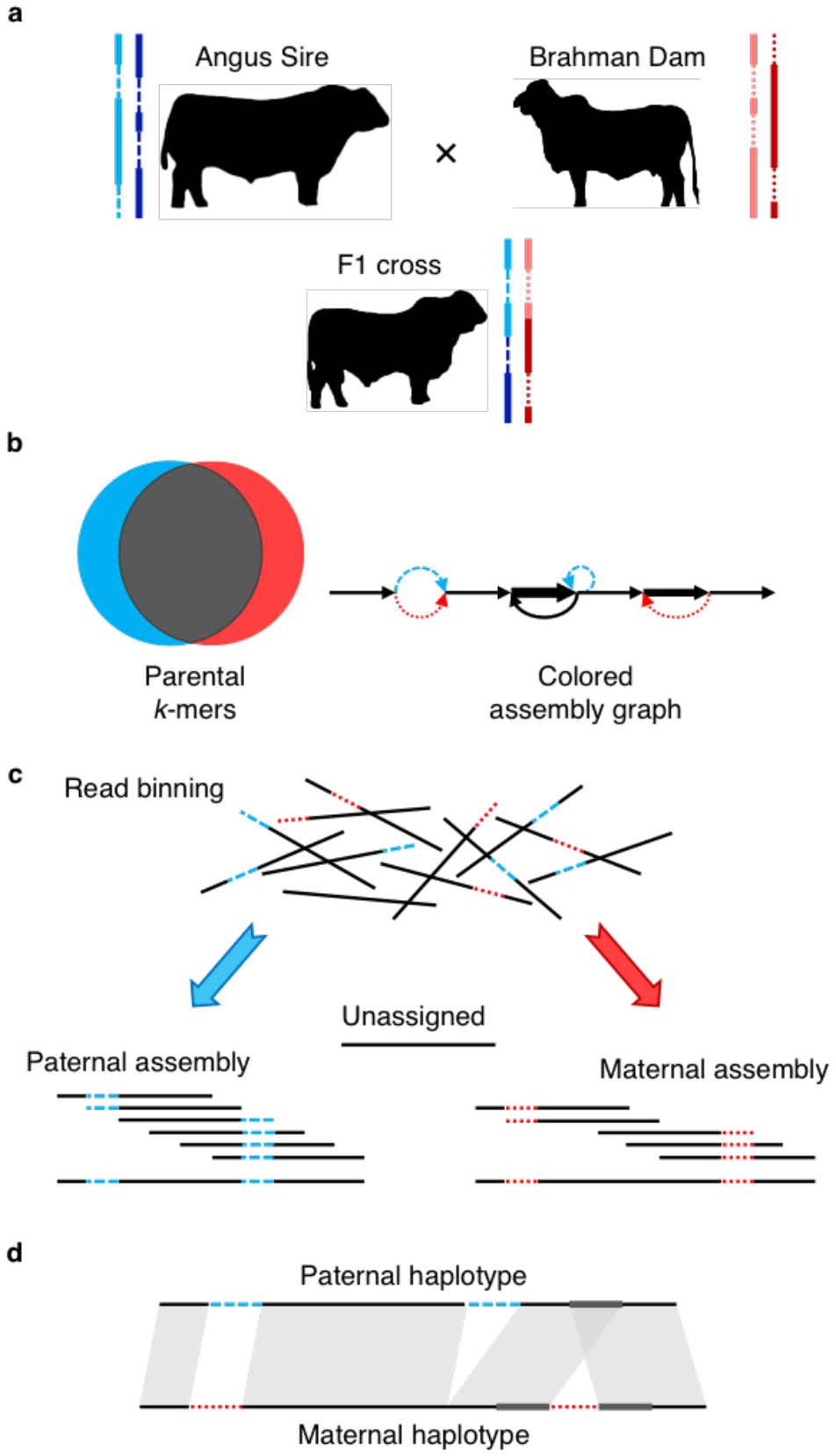
Outline of trio binning and haplotype assembly. a) Two parents constitute four haplotypes including shared sequence in both parents (solid lines) and sequence unique to one parent (dashed lines). The offspring inherits a recombined haplotype from each parent (blue, paternal; red, maternal). b) Short-read sequencing of the parents identifies unique length-*k* subsequences (*k*-mers), which can be used to infer the origin of heterozygous alleles in the offspring’s diploid genome. c) Trio binning simplifies assembly by first partitioning long reads from the offspring into paternal and maternal sets based on these *k*-mers. Each haplotype is then assembled separately without the interference of heterozygous variants. Unassignable reads are homozygous and can be assigned to both sets or assembled separately. d) The resulting assemblies represent genome-scale haplotypes, and accurately recover both point and structural variation.

Trio binning performs best for a uniformly heterozygous offspring, which maximizes the probability that any given read will contain at least one haplotype-specific *k*-mer. Each heterozygous single-nucleotide variant is expected to induce 2*k* haplotype-specific *k*-mers. As a result, the fraction of haplotype-specific *k*-mers is greater than the single-nucleotide heterozygosity. In human, for example, where single-nucleotide heterozygosity is estimated to be only ~0.1%, nearly 2% of the 21-mers are haplotype specific (**Table 1**, **Supplementary Table 1**). Thus, *k*-mers are powerful haplotype markers that can also capture complex insertions, deletions, and fusion events.

**Table 1.**
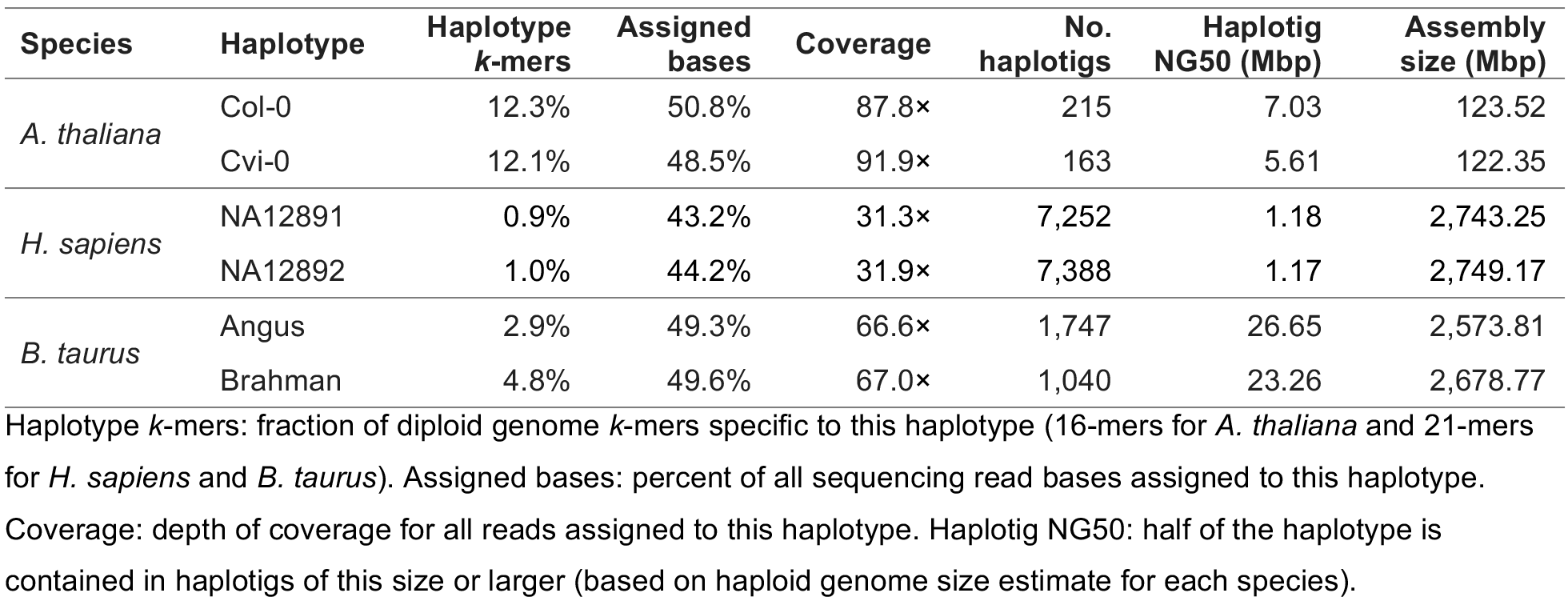
Trio heterozygosity, binning, and assembly statistics.

| Species | Haplotype | Haplotype *k*-mers | Assigned bases | Coverage | No. haplotigs | Haplotig NG50 (Mbp) | Assembly size (Mbp) |
| --- | --- | --- | --- | --- | --- | --- | --- |
| *A. thaliana* | Col-0 | 12.3% | 50.8% | 87.8× | 215 | 7.03 | 123.52 |
|  | Cvi-0 | 12.1% | 48.5% | 91.9× | 163 | 5.61 | 122.35 |
| *H. sapiens* | NA12891 | 0.9% | 43.2% | 31.3× | 7,252 | 1.18 | 2,743.25 |
|  | NA12892 | 1.0% | 44.2% | 31.9× | 7,388 | 1.17 | 2,749.17 |
| *B. taurus* | Angus | 2.9% | 49.3% | 66.6× | 1,747 | 26.65 | 2,573.81 |
|  | Brahman | 4.8% | 49.6% | 67.0× | 1,040 | 23.26 | 2,678.77 |

Haplotype *k*-mers: fraction of diploid genome *k*-mers specific to this haplotype (16-mers for *A. thaliana* and 21-mers for *H. sapiens* and *B. taurus*). Assigned bases: percent of all sequencing read bases assigned to this haplotype.
Coverage: depth of coverage for all reads assigned to this haplotype. Haplotig NG50: half of the haplotype is contained in haplotigs of this size or larger (based on haploid genome size estimate for each species).

Read classification accuracy depends not only on the zygosity of the offspring, but also on sequencing read length and error rate (**Figure 2**). Due to the high error rates in current long-read technologies, the *k-*mer size is also important. It must be long enough to be unique in the genome but short enough that it will not be corrupted by sequencing errors (e.g. *k*≈21 for a 3 Gbp genome). Given current long-read sequencing characteristics (read N50 >15 kb and read accuracy >85%), it is possible to bin and assemble nearly all of a human genome. A small fraction of reads will remain unclassified, but in the three datasets analyzed here, these reads were typically short and derived from either homozygous alleles or identically heterozygous alleles (i.e. both parents share the same heterozygous genotype). The former reads, being homozygous, can be co-assembled with both haplotype bins, while the latter are a limitation of trios and would require additional linkage information to be assigned correctly. However, current read lengths typically exceed the size of such alleles and unclassifiable reads are rare in practice.

**Figure 2.**
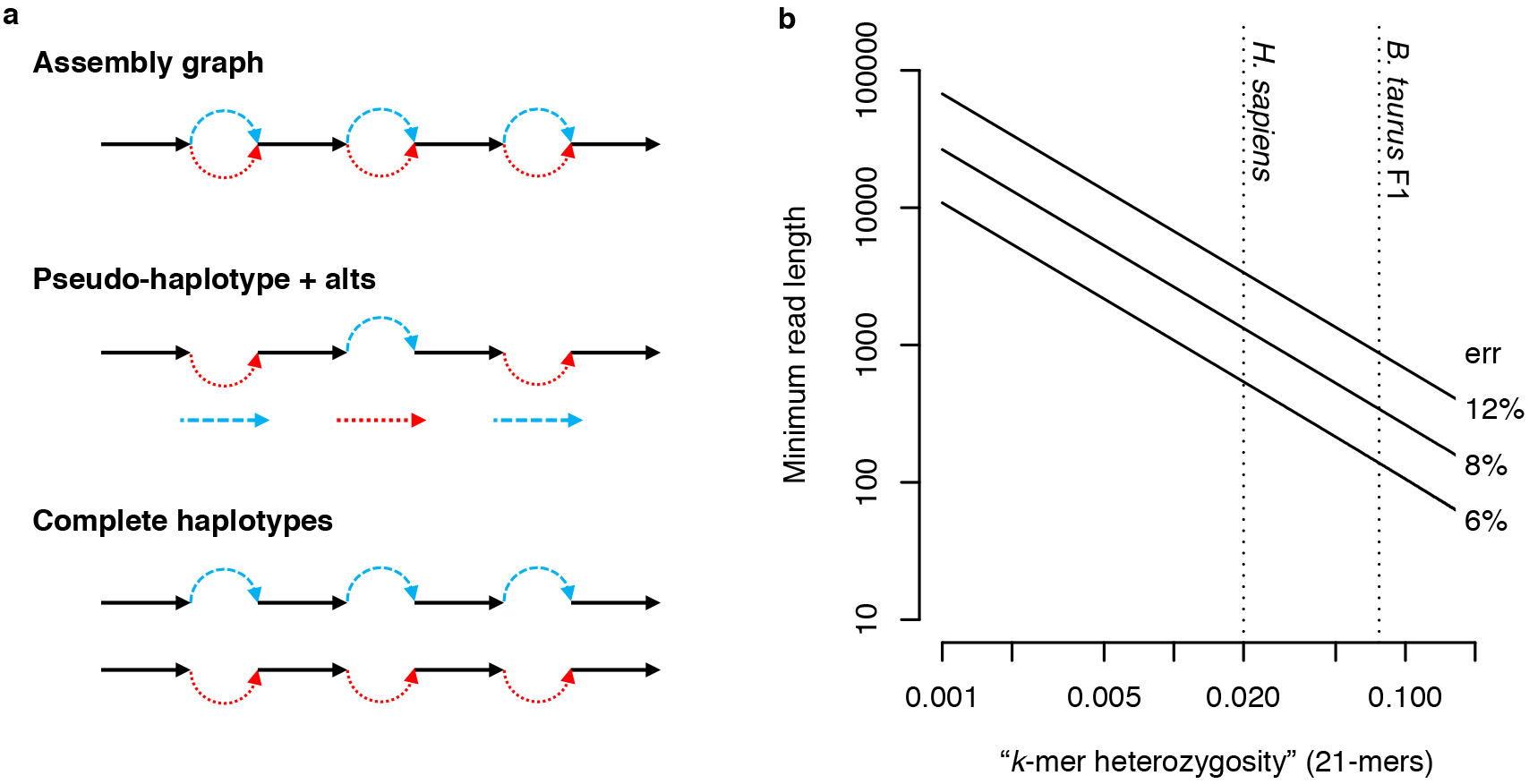
Effect of data characteristics on trio binning. a) Diploid assembly representations shown with homozygous alleles in black and heterozygous alleles (called “bubbles”) colored by haplotype. Graphical representations typically collapse homozygous alleles into a single sequence. A pseudo-haplotype is a path through the diploid graph that separates heterozygous alleles but does not preserve phase between loci. Complete haplotypes represent all alleles and preserve phase across the entire genome. Ability to assign sequencing reads to a haplotype depends on the zygosity of the genome, the sequencing read length, and the sequencing error rate. b) Log-log plot of minimum required read length (y-axis) such that there is a 99% probability of observing at least one haplotype-specific 21-mer per read (negative binomial distribution, Methods), dependent on the sequencing error rate (labels) and fraction of haplotype-specific 21-mers in the genome (x-axis). Dotted vertical lines mark the fraction of heterozygous 21-mers for *H. sapiens* and the *B. taurus* F1 cross.

### Validation on an Arabidopsis cross

The published description of FALCON-Unzip provided a valuable dataset for benchmarking diploid assembly algorithms ^18^. The authors crossed two well-characterized strains of *Arabidopsis thaliana*, Col-0 and Cvi-0, and generated both long-read PacBio and short-read Illumina sequencing reads for the F1 hybrid. Because the parental strains are both highly inbred, recombination is inconsequential and the F1 haplotypes are expected to match the parental genomes, providing a truth set for validation. No short-read data was available for the parental lines, so we inferred haplotype-specific *k*-mers directly from the assemblies. The heterozygosity was estimated to be 1.36% ^28^, or one variant every 73 bases, representing a best-case scenario for diploid assembly.

TrioCanu successfully classified the *A. thaliana* F1 reads by haplotype, resulting in unimodal *k*-mer distributions for the read bins, and an assembly that fully resolved both parental haplotypes (Figure 3, **Supplementary Note 1 and Supplementary Figure 1**). In contrast, rather than reporting complete haplotypes, FALCON-Unzip produces pseudo-haplotypes (“primary contigs”) along with a set of alternate alleles (“associated haplotigs”) that represent haplotype variants. Thus, homozygous alleles are only represented once and there can be considerable haplotype switching within the pseudo-haplotypes. To compare these results to the TrioCanu haplotigs, the FALCON-Unzip primary contigs and associated haplotigs were aligned to the parental genomes to infer the correct mapping between assembly and haplotype.

**Figure 3.**
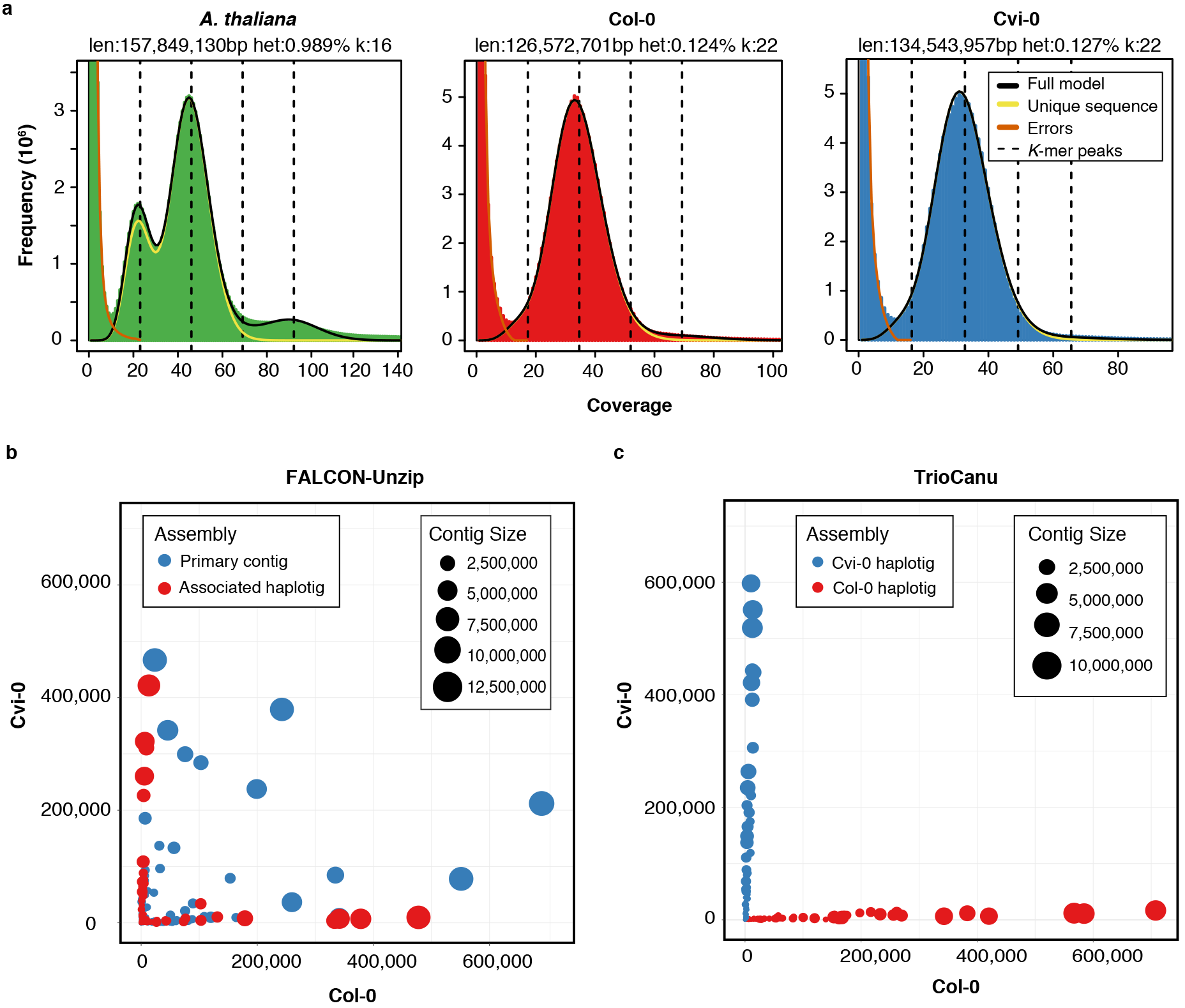
Read and assembly *k*-mer statistics for an *Arabidopsis thaliana* F1 hybrid. a) GenomeScope ^28^ *k*-mer count distributions for the F1 Illumina data (left) and F1 PacBio reads partitioned by haplotype and corrected by TrioCanu (Col-0 and Cvi-0). The Illumina *k*-mer distribution shows two clear peaks, characteristic of a diploid read set, with the 1/2 coverage peak corresponding to haplotype-specific *k*-mers. In comparison, the binned PacBio data shows a normal *k*-mer count distribution, characteristic of a haploid read set. b) Counts of Col-0 (x-axis) and Cvi-0 (y-axis) haplotype-specific *k*-mers in FALCON-Unzip and TrioCanu contigs (colored circles). FALCON-Unzip primary contigs switch between haplotypes, resulting in a mix of *k*-mers from both parents, whereas the FALCON-Unzip associated haplotigs (alts) are smaller, but phase preserving. However, without parental information, these haplotigs cannot be further grouped by haplotype. In comparison, TrioCanu haplotigs contain sequence from only a single haplotype and are automatically sorted into two complete haplotypes.

The TrioCanu haplotypes covered 99.50% and 99.00% of the Col-0 and Cvi-0 parental genomes, respectively, and the alignment identity for both was 99.97%. This exceeds the FALCON-Unzip result of 98.47% and 98.53% coverage and 99.94% and 99.92% identity. The NG50 size of the TrioCanu F1 haplotigs was 7.0 Mbp and 5.6 Mbp for the Col-0 and Cvi-0 haplotypes, respectively, compared to 7.4 Mbp and 6.1 Mbp for the inbred parental genomes (NG50 such that 50% of the haploid genome is contained in haplotigs of this size or greater). Thus, each haplotype of the diploid TrioCanu assembly of the F1 is of comparable quality to the haploid parental assemblies. In comparison, the NG50 of the FALCON-Unzip pseudo-haplotype was longer than TrioCanu at 8.0 Mbp, but contained substantial haplotype switching, as expected. While convenient for some applications, pseudo-haplotypes must be split prior to reconstructing complete haplotypes with the assistance of additional linkage information. This is a difficult and error-prone process. In contrast, TrioCanu haplotigs contain no switching and are inherently partitioned by haplotype.

Accurate structural variant detection from complete haplotypes simply requires a whole-genome alignment of the two. To demonstrate this approach and the accuracy of the TrioCanu haplotypes, we used Nucmer ^29^ and Assemblytics ^30^ to identify a total of 4,828 structural variants (SVs) between the Col-0 reference genome (TAIR10 ^31^) and a *de novo* assembly of the Cvi-0 genome ^18^. Assemblytics classifies SVs in the range ≥50 bp and <10 kbp. These SVs were then compared to those identified by aligning the TrioCanu and FALCON-Unzip F1 assemblies against the Col-0 reference genome. Positive predictive value (PPV) and sensitivity for the TrioCanu Cvi-0 haplotype was 99.1% and 99.2%, respectively, compared to 96.99% and 98.81% for the FALCON-Unzip assembly (primary contigs plus associated haplotigs). However, the FALCON-Unzip PPV is artificially high in this case because variants are only being discovered on one haplotype (i.e. the other haplotype matches the reference and induces no variants). As expected, the TrioCanu F1 Col-0 haplotype showed good agreement with the Col-0 reference genome, differing on average by less than 2 variants per 10,000 bases and 108 SVs, which could represent errors in the assembly, errors in the reference, or true intra-strain variation. NA50s for the TrioCanu and FALCON-Unzip assemblies versus the Col-0 reference genome were 2.4 Mbp and 1.7 Mbp, respectively (NA50 such that 50% of the genome is covered by continuous alignments of this size or greater ^32^).

### A personal, diploid human genome

We next evaluated trio binning on a human trio of European descent (father: NA12891, mother: NA12892, and daughter: NA12878 ^23^), and compared against a Supernova 10x Genomics (linked-read) assembly of NA12878 ^19^. Due to historical population bottlenecks, human genomes typically have a heterozygosity of ~0.1%, which was confirmed for NA12878 via *k*-mer analysis (**Supplementary Figure 2**). This presents a challenge for haplotype recovery, because heterozygous variants are sparse (1 per 1,000 bases on average) and long-range linking information is required to preserve phase. A trio-based approach overcomes this problem because variants can simply be associated with the parent from which they were inherited, preserving phase across the entire genome.

A TrioCanu assembly of NA12878 from 72× PacBio coverage produced a haplotig NG50 of 1.2 Mbp and an assembly size of 2.7 Gbp for each parental haplotype (Table 1 **and Supplementary Note 1**). By comparison, the Supernova assembly from 41× linked-read coverage had a smaller contig NG50 of 103 kbp and NG50 phase block of 4.2 Mbp. Because TrioCanu generates complete haplotypes, the entire genome is in phase and all haplotigs are assigned to the parent from which they were inherited (**Supplementary Figure 3**). For example, the TrioCanu paternal haplotype correctly assembled a known *CYP2C19* substitution ^25^. Comparing the two TrioCanu NA12878 haplotypes using Assemblytics yielded 6,674 structural variants affecting 3.4 Mbp of the genome, including 12 inversions with an average size of 19 kbp. The alignment included 2.67 million single-nucleotide substitutions, matching the expected heterozygosity. Insertions and deletions (indels) between the haplotypes were well balanced, with an enrichment for 300 bp and 6 kbp events, corresponding to human Alu and LINE elements, respectively (**Figure 4a**).

**Figure 4.**
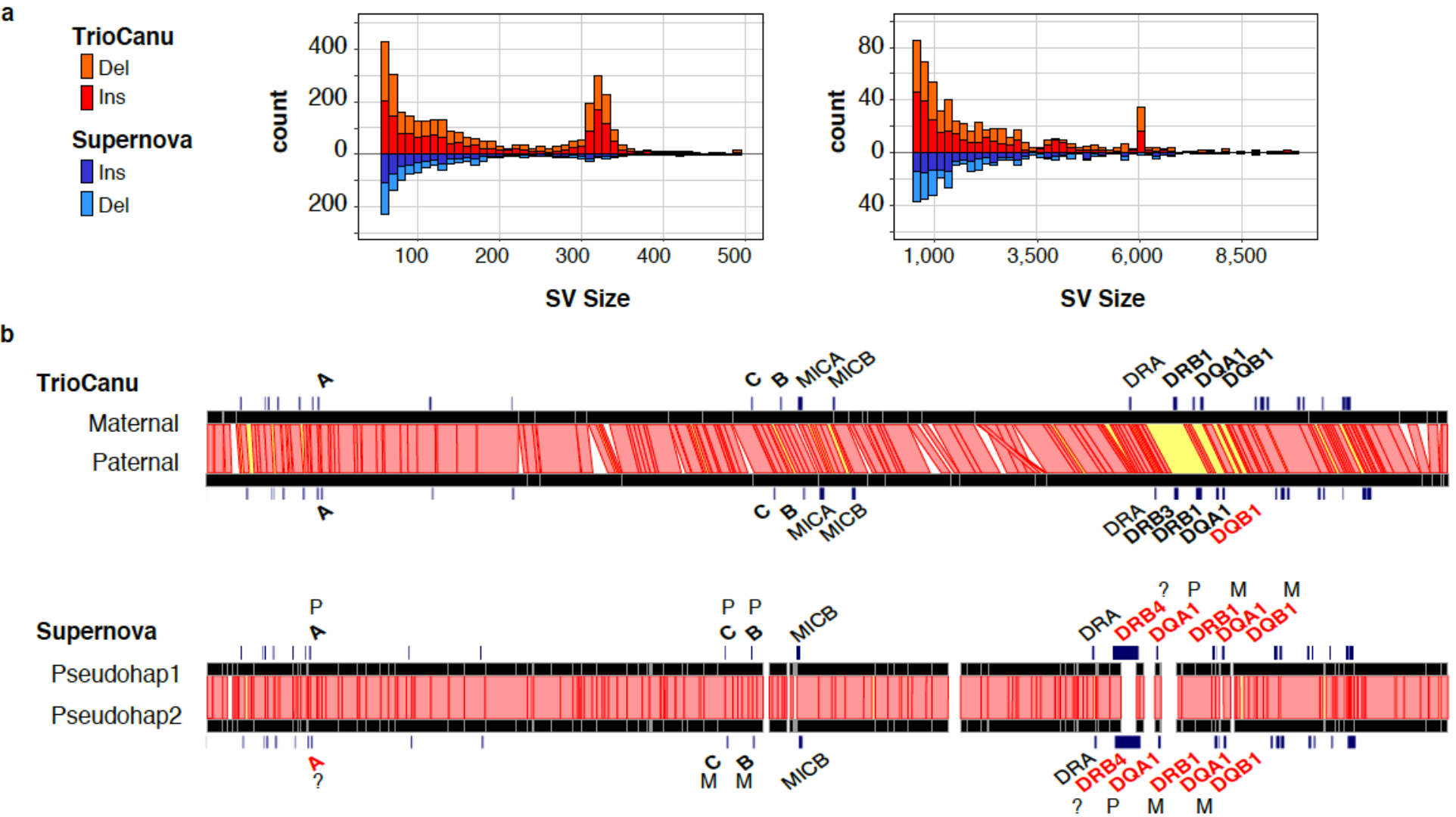
Haplotype variation in a diploid human genome. a) Counts of structural variants between NA12878 haplotypes across the entire genome as reported by Assemblytics ^30^. Canu haplotypes (top, red) showed a balance of insertions and deletions, with peaks at ~300 bp and ~6 kbp corresponding to human Alu and LINE elements, respectively. In comparison, the Supernova pseudo-haplotypes (bottom, blue) were missing these larger structural variants. b) Ribbon visualization ^51^ of MHC haplotypes for human reference sample NA12878 as assembled by TrioCanu from PacBio data (top) and Supernova from 10X Genomics data (bottom). Red bands indicate >95% identity between haplotypes; yellow bands <95% identity; and unaligned in white (gaps and indels). Genes are annotated in black if matching the known truth without error. TrioCanu captured more haplotype variation than Supernova, especially in the highly variable MHC class II region, which contains a long stretch of high sequence divergence (yellow). In addition to phasing the entire region, TrioCanu perfectly reconstructed all typed MHC genes on both haplotypes, with the exception of the paternal DQB1, which contained a single base indel (**Supplementary Table 2**). Supernova produced an overly homozygous reconstruction that incorrectly assembled a majority of genes and introduced false gene duplications (**Supplementary Table 3**).

To measure accuracy, we compared individual SNPs extracted from the TrioCanu and Supernova assemblies against a gold standard variant call set for NA12878 ^25^. Considering only genomic positions covered in both assemblies, sensitivity of TrioCanu was 91.2% versus 90.9% for Supernova and the PPV was 90.2% vs. 93.4%. The lower TrioCanu PPV is likely due to residual consensus errors in the PacBio assembly. A *k*-mer analysis also showed that the TrioCanu assembly is missing some homozygous alleles due to assembly gaps and/or sequencing errors (**Supplementary Figure 4**). A higher coverage of long reads, so that each haplotype approaches 50× coverage, could be expected to reduce both consensus errors and missing alleles. Despite the 10x Genomics assembly having longer input fragments than PacBio (mean 51 kbp vs. 12 kbp), the NG50 perfect phase block for Supernova was 4.3 Mbp vs. 5.6 Mbp for TrioCanu. The few TrioCanu phase errors originate from regions where both parents have identical heterozygous genotypes, which cannot be resolved by the trio method alone without longer read lengths (**Supplementary Note 2**).

The TrioCanu assembly was more structurally accurate than the Supernova assembly. In particular, Supernova missed many larger variants, and assembled fewer Alu and LINE indels relative to TrioCanu (**Figure 4a**). To better understand the structural accuracy of these assemblies, we examined the Major Histocompatibility Complex (MHC), which is a highly repetitive and heterozygous region of the genome that presents a serious challenge for *de novo* assembly. This region contains the human leukocyte antigen (HLA) genes, which have been well characterized for NA12878 ^33^ and serve as a quality check. Supernova did not accurately assemble either MHC haplotype, failed to capture an *HLA-DRB3* gene insertion in the paternal haplotype, and incorrectly reported the majority of the MHC class II region as homozygous (**Figure 4b**). By comparison, TrioCanu correctly assembled both MHC haplotypes, as demonstrated by perfect HLA typing results and only a single base error in the typing genes (**Supplementary Tables 2-3 and Supplementary Note 2**).

### Reference assembly of two cattle breeds using an F1 hybrid

Using trio binning, we sought to generate high-quality, breed-specific reference genomes for Angus and Brahman cattle (examples of the *Bos taurus taurus* and *Bos taurus indicus* subspecies, respectively). We collected ~60× Illumina coverage each for an Angus bull and a Brahman cow, and 134× PacBio coverage in reads ≥1 kbp for their male F1 offspring. Heterozygosity of the F1 was estimated to be 0.9% (**Supplementary Figure 5**).

TrioCanu successfully resolved both F1 haplotypes with a haplotig NG50 exceeding 20 Mbp for each haplotype (NG50: Angus 26.6 Mbp, Brahman 23.3 Mbp) (**Supplementary Note 1**). This far surpasses previous *B. taurus taurus* ^34^ and *B. taurus indicus* ^35^ reference genomes, both of which have contig NG50s <100 kbp. The TrioCanu haplotype assemblies were also more continuous than a Canu assembly of the unbinned data due to heterozygous branching in the assembly graph (NG50: 15.6 Mbp). A FALCON-Unzip assembly of the combined data achieved an impressive NG50 of 31.4 Mbp for the primary contigs, but with the expected pseudo-haplotype switch errors (**Supplementary Figure 6**). Further analysis of *k*-mer distributions in the FALCON-Unzip assembly show less complete haplotype separation, with more homozygous *k*-mers (expected to by 2-copy) occurring either as 1-copy (over-collapsed) or >2-copy (over-split) (Figure 5a, Figure 5b, **Supplementary Figures 7**). For example, FALCON-Unzip over-collapsed roughly twice as many *k*-mers as TrioCanu (**Supplementary Table 4**). Each TrioCanu haplotype was polished using only the haplotype-assigned PacBio reads, and the quality of the final assembly was estimated to be QV47 (accuracy 99.998%, **Supplementary Note 2**), supporting our contention that higher coverage can overcome the limitations of PPV and missing homozygous alleles observed for the lower-coverage human sample. In addition, polishing only the Brahman haplotype using reads from both haplotypes increased the total number of errors more than twofold, despite the increased coverage, due to artifacts introduced by the Angus haplotype. This highlights the advantage of binning reads by haplotype for accurate consensus generation (**Supplementary Note 2**).

**Figure 5.**
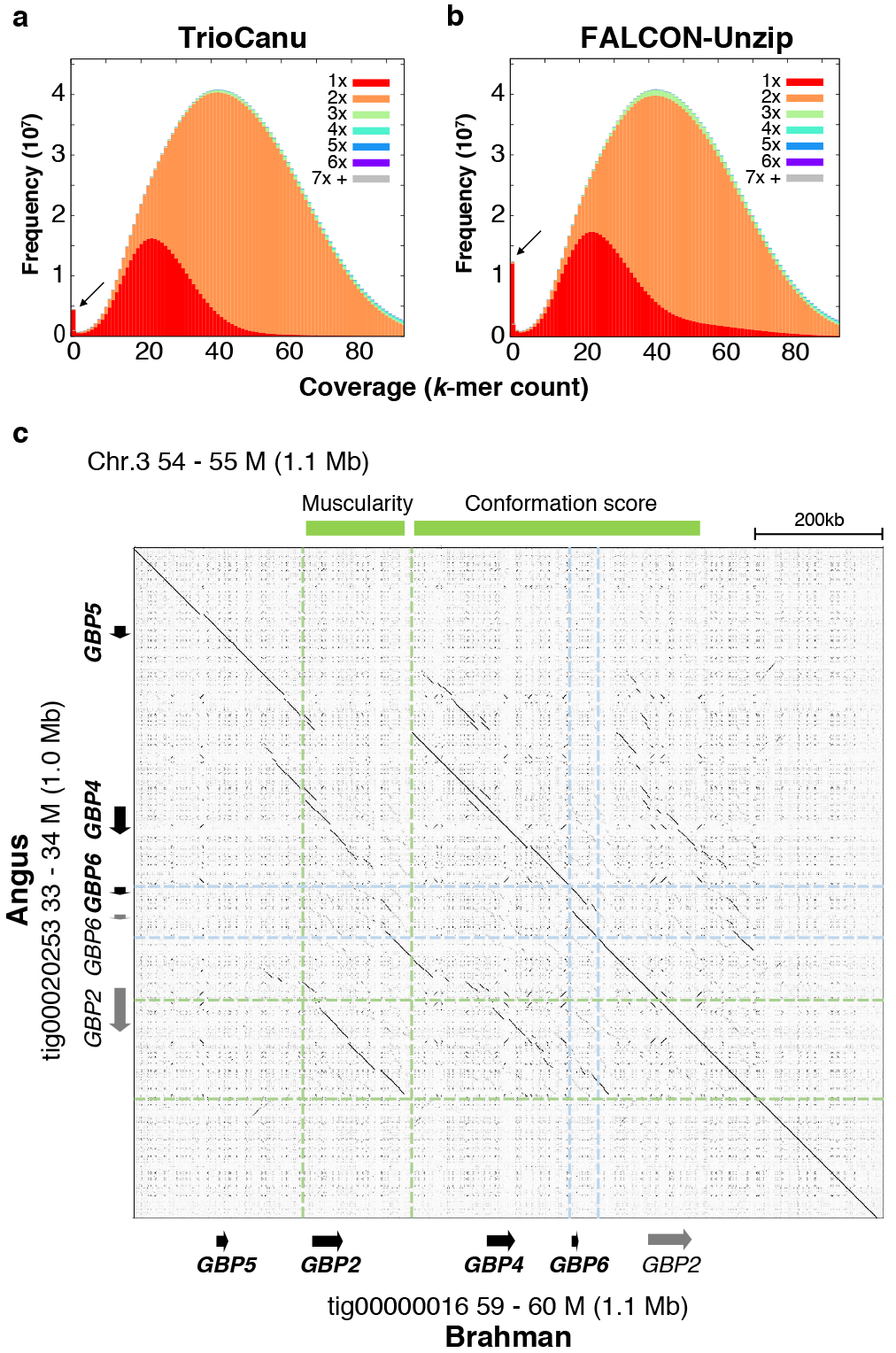
Diploid assembly of a *Bos taurus* F1 hybrid. Stacked *k*-mer histograms from KAT ^52^ comparing a) TrioCanu and b) FALCON-Unzip *k*-mer counts to an independent Illumina dataset of the same individual. The x-axis bins are *k*-mer coverage in the Illumina dataset, and the y-axis is the frequency of those *k*-mers in the Illumina set colored by copy number in the assembly. The FALCON-Unzip distribution has more *k*-mers that do not appear in the Illumina data (arrows), a longer tail of 1-copy *k*-mers (red, collapsed haplotype), and slightly more 3-copy *k*-mers (green, duplicated haplotype). c) Alignment dotplot of the TrioCanu Angus and Brahman haplotypes in a highly heterozygous region containing multiple guanylate binding protein (GBP) genes. Relative to Brahman, the Angus haplotype is missing a ~140 kbp region containing *GBP2*, previously reported to be associated with muscularity (light green). The Angus haplotype also has a duplicated *GBP6*-like sequence (light blue) in a region associated with conformation score (genes marked in grey are highly divergent from known transcripts).

The Angus and Brahman haplotypes aligned to one another with 99.35% identity, and contained 25,245 haplotype-specific structural variants and 124 inversion breakpoints. Common SV sizes corresponded to known retrotransposon families in the *Bos taurus* lineage, including the three most common elements: tRNA-Core-RTE (214 bp), RTE-BovB (1,650 bp), and L1 (5,981 bp) (**Supplementary Figure 8**). One of the most heterozygous regions between the two haplotypes contained notable copy number variations (CNVs) of *GBP2~GBP6* (**Figure 5c**). Interestingly, the Angus haplotype has a large (~140 kb) deletion containing *GBP2*, while Brahman includes a complete version of *GBP2* transcript variant X8. In addition, *GBP6* is partially duplicated only on the Angus haplotype. These regions overlap with a previously reported association of quantitative traits for muscularity and visual conformation scores ^36^, suggesting our breed-specific haplotypes will be important for understanding growth traits in cattle.

BUSCO ^37^ reported 92.6% and 93.4% complete universal single-copy orthologs and a low rate of duplication (1.0% and 1.1%) for the TrioCanu Angus and Brahman haplotigs, respectively, which is consistent with 93.7% completeness and 1.3% duplication for the current *B. taurus taurus* Hereford UMD3.1.1 reference ^34^. To further measure accuracy of the assemblies, we aligned the probe sequence for 735,636 autosomal markers from Illumina’s BovineHD BeadChip to both haplotypes. Only 333 marker loci did not align to either of the TrioCanu haplotypes, and 2,726 and 3,718 were absent from Angus and Brahman, respectively. The 333 marker loci missing from both haplotypes also had low evidence in the parental Illumina data, suggesting that their absence is real in the parental genotypes and not due to incomplete assembly (**Supplementary Figure 9**). The majority of marker sequences missing in one haplotype were also depleted in the corresponding parent’s short read data, but not the other parent, indicating these are haplotype-specific loci correctly phased by the assembly (**Supplementary Figures 10-11**). Switch error between the haplotypes was roughly estimated at 0.68% using independent Hi-C data from the F1 (**Supplementary Note 2 and Supplementary Table 5**).

The Angus and Brahman haplotype assemblies cover 94.2% and 96.2% of the UMD3.1.1 reference genome, respectively, with the Brahman dam haplotype containing the X chromosome and mitochondrial genome (**Supplementary Figures 12-13**). Surprisingly, we identified 3,178 inversions shared by both haplotypes with respect to the reference (mean 9,447 bp, median 4,385 bp, **Supplementary Table 6**). Most of these inversions (94.5%) corresponded with reference scaffold gaps, and the inverted sequences were fully contained within TrioCanu haplotigs. To validate the TrioCanu reconstruction, we used NGM-LR ^38^ and Sniffles ^38^ to identify structural variants in the combined F1 PacBio read set versus the UMD3.1.1 reference assembly and both TrioCanu haplotypes. This identified 3,354 inversions in the UMD3.1.1 assembly, versus just 11 and 20 the Angus and Brahman haplotigs, respectively. Thus, it appears the current cattle reference genome contains systematic inversion errors within its scaffolds (**Supplementary Figures 12-13**). Sniffles also identified over 4-fold more SVs in the UMD3.1.1 assembly versus our haplotigs and 1.8-fold more deletions than insertions (**Supplementary Table 6**), which was also evident from the Assemblytics output (**Supplementary Figures 14-15**), suggesting artifactual duplications in the UMD3.1.1 assembly. Comparison against a new long-read reference sequence ARS-UCDv1.0.11 (B. Rosen, personal communication) of the same Hereford animal used for UMD3.1.1, showed no apparent indel bias and returned 3-fold and 2-fold fewer variants versus the Angus and Brahman assemblies, respectively, further supporting error in the UMD3.1.1 assembly (**Supplementary Figures 16-17**). A comparison between our Angus and Brahman haplotypes mirrored a comparison between ARS-UCDv1.0.11 and the Brahman haplotype (**Supplementary Figure 8**). In contrast, the existing short-read *B. taurus indicus* genome contains few variants over 500 bp (**Supplementary Figure 18**), and likely inherits assembly errors from UMD3.1.1 due to use of a reference-guided assembly approach ^35^ (**Supplementary Figure 19**).

## Discussion

Technologies that produce sequencing reads longer than common repeat lengths have dramatically improved the continuity of genome assemblies, but accurate reconstruction of diploid and polyploid genomes has remained a challenge. As a result, most genome projects select an inbred individual for sequencing or settle for a mosaic assembly that represents an artificial mixture of haplotypes. This can introduce significant errors in the downstream analyses. Furthermore, accurate representation of haplotypes is essential for studies of intraspecific variation, chromosome evolution, and allele-specific expression.

Here we have demonstrated that trio binning facilitates complete haplotype assembly for heterozygous diploid genomes, including human. This strategy has several advantages over traditional approaches. First, trio binning recovers the true haplotypes of a viable organism, and requires fewer resources than inbreeding. It is also applicable to organisms that have long generation times or are otherwise recalcitrant to inbreeding. Second, by isolating haplotype variation prior to assembly, the resulting assembly graphs are simplified. As a result, haplotype-specific assemblies can exceed the continuity of merged diploid assemblies. Third, our approach is able to accurately reconstruct structurally heterozygous alleles that can be important factors in adaptation and immunity (e.g. the MHC) and have previously been linked to quantitative traits (e.g. GBP genes). We have shown that such sequences are often mis-assembled by alternative approaches.

Trio binning is fully compatible with downstream analysis tasks. Unlike graph-based assembly representations, which require a specialized bioinformatics toolchain, linear haplotypes can be easily analyzed with existing methods. For example, the partitioned read sets can be reused for haplotype-specific gap filling ^39^ and consensus polishing ^40, 41^, and we have shown that polishing with haplotype-specific reads achieves a more accurate consensus sequence. Given sufficient haplotype divergence and read lengths, nearly all reads are assigned to the correct haplotype. However, for genomes with lower heterozygosity, long homozygous alleles may receive lower coverage and quality due to a lack of assigned reads. In these cases, homozygous reads can be assigned to both haplotypes to boost coverage at the risk of masking some true variants. Additional processing after assembly could correct for this, for example, by mapping the parental short read data to identify missed variants and correct switch error. Alternatively, the accuracy of long-read binning could be improved by more sophisticated classification (e.g. using spaced *k*-mers ^42^) or the integration of additional data types (e.g. Hi-C). The latter option may allow partial haplotype binning without the use of a trio.

Long-read trio binning, as described here, is the first method able to assemble complete haplotypes from a heterozygous genome, and has immediate applications to reference genome construction as well as human and agricultural genomics. New reference genomes will benefit from the improved assembly accuracy and continuity of this approach. For agricultural genomics, trio binning can be used to study breed diversity and has the advantage of producing two reference-quality haplotypes from a single individual. Our assembly of an outbred F1 resulting from a cross between Angus and Brahman cattle produced two breed-specific haplotypes that improve upon and correct the current best reference genomes for both subspecies. These haplotype-specific reference sequences provide an important resource for understanding genetic variation in cattle. The more general idea of haplotype binning should also work well for polyploid plant genomes (e.g. bread wheat) by utilizing species markers (rather than parental markers) to pre-partition reads by haplotype. For human genomics, our approach is a viable method for reconstructing complete, personalized haplotypes, and could be used to generate a more complete database of human haplotype variation.

## Methods

### Data and code availability

Sequencing data for the cattle trio is available under NCBI BioProject PRJNA432857.
All other sequencing data was obtained from public sources. Data accessions, software versions, and commands used to produce the described results are provided in **Supplementary Note 1**. Assembly files, accession numbers, and other miscellaneous information can be found at: https://gembox.cbcb.umd.edu/triobinning/index.html. Prototype code used to build *k*-mer sets, subtract parental *k*-mers, and classify reads is available from https://github.com/skoren/triobinningScripts. Trio binning will be fully incorporated into Canu starting with release v1.7.

### Haplotype *k*-mer identification

TrioCanu automates *k*-mer counting, thresholding, and set operations to identify haplotype-specific *k*-mers. All *k*-mers are counted using Meryl, a sort-based *k*-mer counter used within Canu that allows linear-time *k*-mer set operations. First, a *k*-mer frequency distribution is obtained by counting *k*-mers in the parental genomes. This distribution is examined to eliminate *k*-mers likely to be erroneous (low copy) or from genomic repeats (high copy), leaving only *k*-mers from unique homozygous or heterozygous genome sequences ^43^. For *k*-mer coverage *x* and frequency *y*, the optimal low coverage threshold is determined by finding the first critical point *y′* = 0 and its corresponding coverage *x*_0_ and frequency *y*_0_. The same frequency cutoff *y*_0_ is used to determine the high coverage threshold *x*_1_. A *k*-mer set *D* for each haplotype is drawn from all haplotype *k*-mers with coverage *x*_0_ < *c* < *x*_1_. For two parental haplotypes *i* and *j*, haplotype-specific *k*-mer sets are then constructed as *H*_*i*_ = *D*_*i*_ − *D*_*j*_ and *H*_*j*_ = *D*_*j*_ − *D*_*i*_.

Smaller *k-*mers are more likely to avoid sequencing error in the reads, so it is preferable to choose a small value for *k*. However, *k* must be large enough to minimize random *k-*mer collisions in the genome. For example, the total space of 16*-*mers is only 4^16^ or 4.29 billion, close to the total number of *k*-mers in a 3 Gbp mammalian genome, increasing the chance that some *k-* mer may occur multiple times simply by chance (and not homology). Given a genome size *G* and tolerable collision rate *p*, an appropriate *k* can be computed as *k* = log_|*∑*|_ (*G*(1 − *p*)/*p*)^44^.

According to this formula, we used *k*=16 for *A. thaliana* and *k*=21 for *H. sapiens* and *B. taurus*.

*A. thaliana*, *H. sapiens*, and *B. taurus* were assembled prior to TrioCanu automation, and haplotype-specific *k*-mer thresholds were identified manually as described in **Supplementary Note 1**. For *A. thaliana*, which was lacking Illumina data for the parents, *k*-mers were collected from assemblies of the parents, excluding repetitive *k*-mers occurring more than 10 times. For *H. sapiens* and *B. taurus*, haplotype-specific *k*-mers were collected from unassembled, short read sequencing of the parents. Low and high *k-*mer coverage thresholds were chosen manually as *x*_0_=30 and *x*_1_=160 for *H. sapiens* and *x*_0_=11 and *x*_1_=100 for *B. taurus*. Retrospective application of the automated thresholding method selected similar thresholds of {[25,143], [27,147]} and {[10,57], [10,67]} for *H. sapiens* and *B. taurus*, respectively.

### Haplotype binning

Haplotype binning is a general strategy for partitioning a read set into haplotype groups prior to assembly. The number of haplotypes is not necessarily limited to two. Given *N* haplotypes, the goal is to identify haplotype-specific *k-*mers that are exclusive to one haplotype. Given a database of haplotype-specific *k-*mers, the number of specific *k-*mers from each haplotype is counted in each read. It is expected that *k*-mers in a single read will be from the same haplotype, but due to sequencing errors it is possible to observe spurious *k*-mers from a different haplotype. Therefore, the observed haplotype-specific *k*-mer counts are normalized by the database size to control for the different *k*-mer set sizes of the parents. Reads are then assigned to the haplotype with the most matching haplotype-specific *k*-mers. In the event of a tie or too few haplotype-specific *k*-mers, the read is marked as ambiguous. Finally, the *N* read bins are passed to Canu for assembly, with the option to include the ambiguous reads in all bins.

Whether a read can be correctly classified is a function of the *k*-mer heterozygosity *h*, read length *l*, read error rate *e*, and *k*-mer size *k*. For simplicity of modeling, errors and haplotype differences are assumed to be random point mutations, and heterozygosity *h* is defined as the fraction of genomic *k*-mers that are haplotype specific. It is assumed that *k* is large enough to avoid chance collisions. A read of length *l* contains *l* − *k* + 1 *k*-mers. The probability of a single *k-*mer surviving uncorrupted is (1 −*e*)^*k*^, and the expected number of uncorrupted *k-*mers in a read is (*l* − *k* + 1)(1 −*e*)^*k*^. The expected number of haplotype-specific *k-*mers in a read is *h*(*l* − *k* + 1), and the number of surviving haplotype specific *k-*mers in a read is *h*(*l* − *k* + 1)(1 −*e*)^*k*^. Thus, for a typical long sequence read with *e*=0.12 and *l*=15,000, and *k*-mer heterozygosity *h*=0.001, the expected number of surviving haplotype-specific 16-mers is 2 and 21-mers is 1. Increasing divergence to *h*=0.01 increases the expected number of 16-mers to 19 and 21-mers to 10.

### Validation

Classification accuracy was evaluated using a truth set of *A. thaliana* parental reads. The simple majority-wins classification heuristic showed a good sensitivity/specificity trade off, exceeding 80% true positive rate with <20% false positive rate (**Supplementary Figure 20**). False positives include homozygous reads which do not affect the resulting assembly, and a small fraction of mis-classified heterozygous reads. These will be outvoted by the majority of correctly classified reads when building the haplotype consensus. If high specificity is required, the classifier can be tuned to require more than a simple majority of haplotype-specific *k*-mers.

Assembly alignments were performed with MUMmer 3.23 ^29^ with the commands

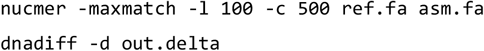

GRCh38 ^45^ excluding ALT loci was used for *H. sapiens.* TAIR10 was used for *A. thaliana*. GCF_000003055.6 with chromosome Y from NC_016145.1 was used for *B. taurus* and AGFL00000000.1 for *B. indicus.* A genome size of 119,667,750 was used for *A. thaliana* (TAIR 10 length), 3,098,794,149 for *H. sapiens* (GRCh38 primary assembly excluding alternates), and 2,713,423,491 for *B. taurus* (the UMD 3.1.1 reference plus the Y chromosome).

Parent-specific *k*-mers were used to estimate switch error within assembly contigs. MHC typing was run as previously described ^46^ with the truth set from Dilthey *et al.* ^47^. *B. taurus* markers used in BovineHD BeadChip (Illumina Inc., San Diego, CA) were used to identify missing regions in the assemblies as well as haplotype-specific sequences. Illumina data was used to estimate QV by mapping with BWA-MEM ^48^ and identifying variants with Freebayes ^49^. Repeats in the *Bos taurus* genome were downloaded from the UCSC genome browser ^50^ (**Supplementary Note 2**).

### Sample preparation and sequencing of the cattle trio

The animals used were part of the Davies Epigenetics and Genetics Resource at the University of Adelaide, Australia, and were established and sampled using procedures approved by the animal ethics committee of the University. A cow of the Brahman breed (subspecies *Bos taurus indicus*) was bred by artificial insemination using semen from a bull of the Angus breed (*Bos taurus taurus*). The Brahman female had been previously typed for mitochondrial DNA haplotype to verify the maternal lineage as indicus-specific. At day 153 post-insemination, the animal was sacrificed and the fetus removed for dissection. The fetal lung was removed immediately into liquid nitrogen, and DNA was extracted using a salting out procedure. Briefly, approximately 100 mg of tissue was ground under liquid nitrogen to a powder, and transferred to a tube containing 2.26 mL of nuclei lysis mixture (2 mL buffer NFB composed of 10 mM Tris-HCL pH 8.0, 0.4 M NaCl, 2 mM EDTA, plus 0.2 mL 10% SDS, plus 0.06 mL 10 mg/mL RNase A). Tissue and solution were mixed by inversion for 2 minutes, then set to shake slowly at 37°C 1 hour. Protein digestion was performed by adding 0.025 mL Proteinase K (20 mg/mL) and returning to the shaker overnight (approximately 16 hours). Protein was removed by addition of 1.25 mL of saturated NaCl, followed by vigorous hand shaking for 15 seconds and centrifugation 2250 × g, 20 minutes, 4 °C. The clarified supernatant was transferred to a tube containing 8 mL of cold 100% ethanol, and DNA was precipitated by gentle rocking of the solution. The DNA was transferred using a glass rod and washed twice in tubes containing 5 mL of 70% ethanol. The pellet was then transferred to a 1.5 mL tube and air dried for 10 minutes at room temperature. DNA was removed from the glass rod by dissolving in 0.25 mL of solution containing 10 mM TrisHCl pH 8.0 and 0.1 mM EDTA overnight at 4 °C. Parental DNA samples were extracted using standard phenol-chloroform based procedures.

Sequence libraries for the parents and the fetus were prepared with TruSeq PCR-free preparation kits as directed by the manufacturer (Illumina, San Diego, CA). The three libraries were sequenced in separate runs, with no other libraries present in the flow cell, on a NextSeq500 instrument using 2×150 paired end reads with High Output Kit v2 chemistry. The libraries employed unique indexes and, despite being in separate runs, only reads with appropriate indexes for the library were used for analysis to prevent any cross-contamination between the sire, dam, or fetal library data.

Libraries for SMRT sequencing were constructed as recommended by the manufacturer (Procedure P/N 100-286-000-07, Pacific Biosciences, Menlo Park, CA), using a 15 kb cutoff for size selection on the BluePippin instrument (Sage Science, Beverly, MA). A total of 12 library preparations were used, nine of which were sequenced using P6/C4 chemistry on an RSII instrument (Pacific Biosciences, Menlo Park, CA) which generated approximately 152 Gb of sequence, and the other three libraries were sequenced on a Sequel instrument which generated another 132 Gb.

## Acknowledgements

We thank William Thompson, Kristen Kuhn, Kelsey McClure, and Robert Lee for technical assistance, and Tina Graves-Lindsay and Washington University in St. Louis for public release of the PacBio NA12878 data. SK, AR, BPW, and AMP were supported by the Intramural Research Program of the National Human Genome Research Institute, National Institutes of Health. SH and JLW are funded from the JS Davies bequest to the University of Adelaide. TPLS was supported by USDA-ARS Project 3040-31000-100-00D. DMB was supported by USDA-ARS Project 5090-31000-026-00-D. This research was also supported by a grant of the Korea Health Technology R&D Project through the Korea Health Industry Development Institute (KHIDI), funded by the Ministry of Health & Welfare, Republic of Korea (grant number: HI17C2098). This work utilized the computational resources of the NIH HPC Biowulf cluster (https://hpc.nih.gov). Mention of trade names or commercial products in this publication is solely for the purpose of providing specific information and does not imply recommendation or endorsement.

## Author contributions

AMP and TPLS conceived and coordinated the project. SK and AR designed the trio-binning method. SK, AR, and BPW implemented the software. SK, AR, BPW, ATD, DMB, SBK, and AMP performed analyses. SH designed and performed breeding experiments and sample collections. JLW contributed to development of the concept and provision of samples. TPLS
performed sequencing. SK, AR, TPLS, JLW, and AMP wrote the manuscript. All authors approved the final manuscript.

